# Geothermal stickleback populations prefer cool water despite multigenerational exposure to a warm environment

**DOI:** 10.1101/615005

**Authors:** Natalie Pilakouta, Shaun S. Killen, Bjarni K. Kristjánsson, Skúli Skúlason, Jan Lindström, Neil B. Metcalfe, Kevin J. Parsons

## Abstract

Given the threat of climate change to biodiversity, a growing number of studies are investigating the potential for organisms to adapt to rising temperatures. Earlier work has predicted that physiological adaptation to climate change will be accompanied by a shift in temperature preferences, but empirical evidence for this is lacking. Here, we test whether exposure to different thermal environments has led to changes in preferred temperatures in the wild. Our study takes advantage of a ‘natural experiment’ in Iceland, where freshwater populations of threespine sticklebacks (*Gasterosteus aculeatus*) are found in waters warmed by geothermal activity year-round (warm habitats), adjacent to populations in ambient-temperature lakes (cold habitats). We used a shuttle-box approach to measure temperature preferences of wild-caught sticklebacks from three warm-cold population pairs. Our prediction was that fish from warm habitats would prefer higher water temperatures than those from cold habitats. We found no support for this, as fish from both warm and cold habitats had an average preferred temperature of 13°C. Thus, our results challenge the assumption that there will be a shift in ectotherm temperature preferences in response to climate change. In addition, since warm-habitat fish can persist at relatively high temperatures despite a lower temperature preference, this suggests that preferred temperature alone may be a poor indicator of a population’s adaptive potential to a novel thermal environment.

## INTRODUCTION

Climate change poses a substantial threat to biodiversity, as rising temperatures are altering abiotic and biotic conditions and, in turn, imposing novel selection pressures on organisms (Crozier & Hutchings 2014). In light of this, there is now a pressing need to understand the capacity of populations to respond and adapt to increasing temperatures (Crozier & Hutchings 2014). Due to their lability, behavioural responses can play a role in facilitating or hindering adaptation to new thermal environments, depending on certain attributes of the species and environmental conditions (Sih et al. 2011, Tuomainen & Candolin 2011, van Jaarsveld et al. 2021).

Thermoregulatory behaviour could be especially important in the context of climate change, as it allows animals to buffer the effects of temperature changes by choosing suitable microhabitats (Fangue et al. 2009, Kearney et al. 2009, Huey et al. 2012, Fey et al. 2019). Animals usually seek temperatures that coincide with their optimal physiological performance and growth, which tends to be determined by their thermal evolutionary history (Jobling 1981, Kellogg & Gift 1983, Diaz et al. 2007, Pörtner & Farrell 2008, Köhler et al. 2011; but see Huey & Bennett 1987, Martin & Huey 2008, Buckley et al. 2022). Thermoregulatory behaviour is particularly important in ectotherms, because ambient temperature directly influences their body temperature, making them vulnerable to temperature changes (Zuo et al. 2011). Thus, it has been proposed that ectotherms should adapt to increasing environmental temperatures through shifts in temperature preferences that accompany evolutionary changes in their physiology (Kearney et al. 2009, Gilbert & Miles 2017, Logan et al. 2018, Catullo et al. 2019). Yet, it is still unclear whether temperature preference has the capacity to evolve in response to long-term changes in the thermal environment (Paranjpe et al. 2013, Logan et al. 2018).

We addressed this by testing whether exposure to a warm environment over multiple generations in the wild leads to the evolution of higher preferred temperatures. We used a unique ‘natural experiment’ in Iceland, where freshwater populations of threespine sticklebacks (*Gasterosteus aculeatus*) are found in waters warmed by geothermal activity (warm habitats) adjacent to populations in ambient-temperature lakes (cold habitats). This study system provides repeated examples of populations experiencing long-term contrasting thermal environments over a small geographic scale, thereby avoiding the confounding factors associated with latitudinal or elevational comparisons. There is evidence for strong divergence in physiology of sticklebacks from these habitats, with warm populations showing a lower standard metabolic rate than cold populations when compared at the same temperature (Pilakouta et al. 2020). In addition, we have demonstrated heritable divergence in morphology, with sticklebacks from warm habitats being more deep-bodied with shorter jaws (Pilakouta et al. *in press*). Warm-habitat sticklebacks also tend to be less social (Pilakouta et al. 2022). Together, the consistency of these changes across populations suggests that they are undergoing adaptative divergence.

Given that standard metabolic rate and temperature preference have previously been shown to be negatively correlated (Killen 2014), we predicted that fish from warm habitats would prefer higher water temperatures than fish from cold habitats. We also expected that this potential divergence in temperature preference may be more pronounced in populations that have been exposed to warm water for more generations. Lastly, we examined individual activity to determine whether inter-individual variation in activity can explain differences in temperature preferences (Killen 2014).

## METHODS

### Study animals

We collected adult threespine sticklebacks from six freshwater populations in Iceland in May–June 2016 (Table 1, Figure 1). Two of these populations were allopatric, meaning that the warm and cold habitats were in neighbouring but separate water bodies with little to no potential for gene flow (Table 1). Because these populations were both located in close proximity to the marine habitat (and were thus more likely to be directly invaded by a common marine ancestor), we considered them a comparable warm-cold ‘population pair’. We also sampled two sympatric warm-cold population pairs, where the warm and cold habitats were in the same water body with no physical barriers between them (Table 1). Warm- and cold-habitat sticklebacks differ in their morphology and physiology both in sympatry and allopatry (Pilakouta et al. 2020, Pilakouta et al. *in press*).

**Figure 1.**
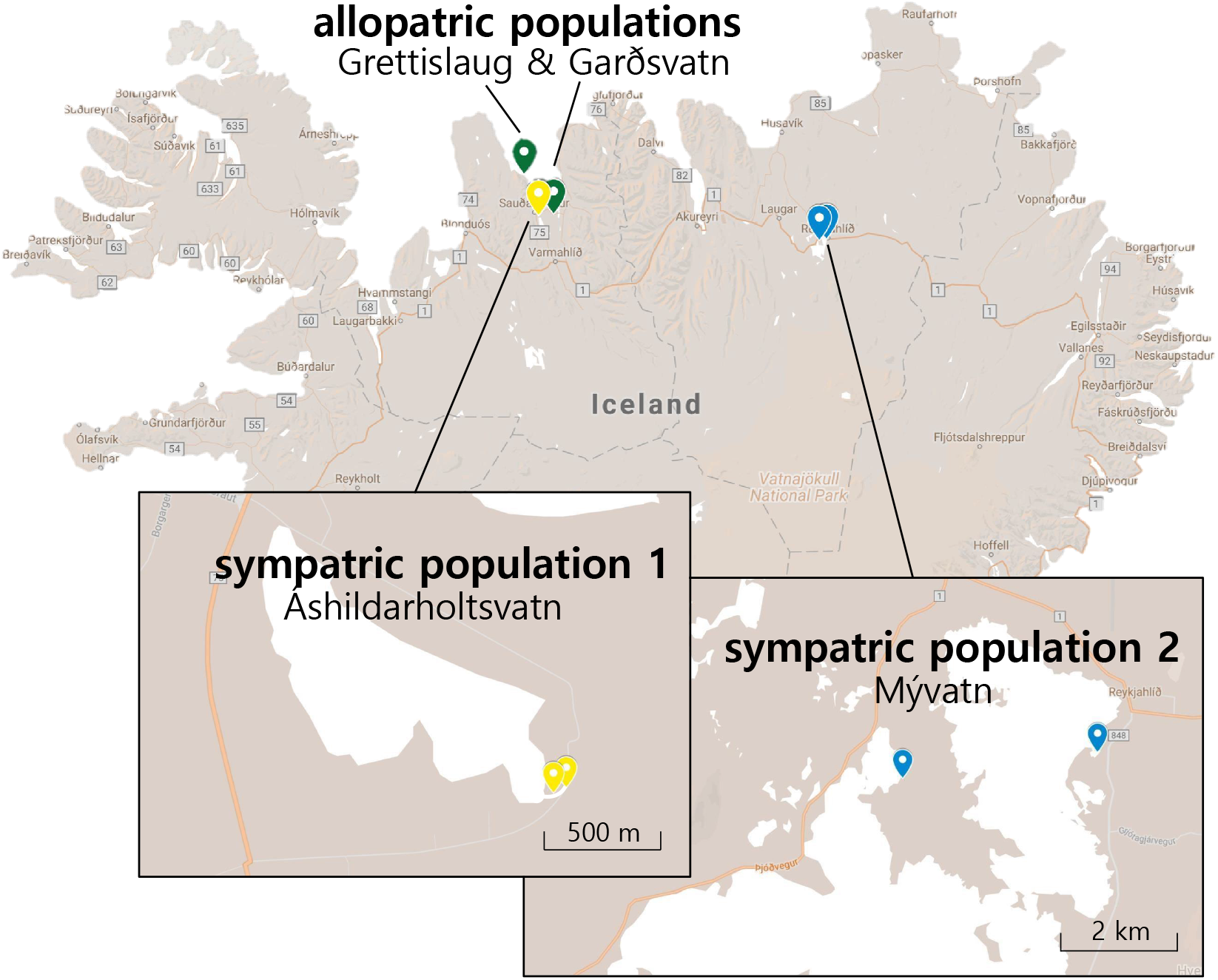
Map of Iceland showing the sampling locations of warm and cold-habitat sticklebacks we collected for this study. Each of the three population pairs is indicated by a different colour.

**Table 1.** Sampling locations of warm- and cold-habitat sticklebacks collected in May–June 2016. Distance refers to how far apart the warm-habitat and cold-habitat sample sites are for each warm-cold pair. All cold habitats have existed since the last glacial period and are therefore approximately 10,000 years old, whereas warm habitats can be classified as either young (<100 years old) or old (>1,000 years old). The summer and winter temperatures listed are the average water temperatures recorded at each sampling location during the summer and winter, respectively.

| Population pair | Water body | Thermal habitat | Age of warm habitat | Distance (km) | Summer temperature (°C) | Winter temperature (°C) | Sample size |
| --- | --- | --- | --- | --- | --- | --- | --- |
| allopatric populations | Grettislaug | warm | old | 21.04 | 24.9 | 13.5 | 10 |
|  | Garðsvatn | cold |  |  | 14.6 | 2.2 | 10 |
| sympatric population 1 | Áshildarholtsvatn | warm | young | 0.05 | 24.1 | 12.5 | 10 |
|  |  | cold |  |  | 12.2 | 3.4 | 10 |
| sympatric population 2 | Mývatn | warm | old | 3.18 | 22.8 | 22.0 | 10 |
|  |  | cold |  |  | 11.5 | 1.0 | 10 |

The cold habitats have likely existed for thousands of years, since the last glacial period (Einarsson et al. 2004), but there is some variation in the age of the warm habitats (Table 1). The ‘Mývatn warm’ and Grettislaug sites have been naturally heated by geothermal activity for over 2000 years (Hight 1965, Einarsson 1982). In contrast, the ‘Áshildarholtsvatn warm’ population originated only 50–70 years ago, fed by excess hot water runoff from nearby residences using geothermal heating. Since the generation time for threespine sticklebacks is about 1 year, the age of the warm habitats corresponds to the maximum number of generations each population pair may have been separated (Table 1). These different timescales made it possible to qualitatively infer whether populations exposed to higher temperatures for a relatively short time have diverged in temperature preference to the same extent as much older populations.

### Animal husbandry

We placed 15 individuals from each population in 10-litre tanks in a common recirculation system. They were fed *ad libitum* twice a day with a mixture of frozen bloodworms, *Mysis* shrimp, and *Daphnia*. Before the experiment, all fish were anaesthetised using benzocaine and marked with visible implant elastomer tags (Northwest Marine Technology Inc) to allow individual identification. They were kept at 15°C and a 12h: 12h daylight cycle for at least one month before being used in the experiment. The acclimation temperature of 15°C was chosen because it is an intermediate temperature for the warm and cold populations (Pilakouta et al. 2020). It is close to the maximum temperature experienced by cold populations in the summer and the minimum temperature experienced by warm populations in the winter (Table 1).

### Experimental set up

We tested each individual’s temperature preference using a classical shuttle-box approach in which the animal is allowed to behaviourally adjust the temperature of its surroundings (McCauley 1977, Schurmann & Steffensen 1992, Westhoff & Rosenberger 2016, Macnaughton et al. 2018). Our shuttle-box apparatus (Loligo Systems) consisted of two circular 40-cm diameter choice chambers joined by a 10-cm passage way (Killen 2014). The left chamber was designated as the warm chamber and the right one as the cold chamber. Each choice chamber was filled with water to a depth of 7 cm and was attached to its own external buffer tank. The set-up also included a heating reservoir, kept at 30°C using aquarium heaters, and a cooling reservoir, kept at 4°C by an external chilling unit. To adjust the temperature within each buffer tank, water was pumped from the buffer tanks through steel coils in the heating and cooling reservoirs. The water temperature in the two choice chambers was continually monitored using in-line temperature probes connected to a computer-driven temperature controller and data acquisition system (DAQ-M, Loligo Systems). In turn, the temperature within each chamber was controlled by software (Shuttlesoft, Loligo Systems), which adjusted the flow from reservoir tanks to change the temperature in the choice chambers as required.

We used two ways of adjusting temperatures in the shuttle-box: a static mode and a dynamic mode. In the static mode, there was a constant temperature in each choice chamber with a 2°C differential between them. In the dynamic mode, the warm and cold chamber temperatures changed depending on the location of the focal fish but always maintained a 2°C differential. When a fish moved into the warm chamber, the temperature increased at a rate of 2°C h ^-1^ in both chambers, whereas when a fish moved into the cold chamber, the temperature decreased at a rate of 2°C h ^-1^ in both chambers. Thus, by moving between the warm and cold chambers in response to changing temperatures, fish could regulate the water temperature they experienced (Schurmann & Steffensen 1992, Killen 2014). Fish movements were tracked using a camera (uEye, Imaging Development Systems GmbH) mounted above the shuttle-box.

### Experimental protocol

All experimental fish were nonbreeding, uninfected adults with their mass ranging from 0.74 to 3.02 g (mean ± SD = 1.82 ± 0.54). Our sample size was *n* = 10 for each population (total *n* = 60). Because feeding history can influence preferred temperature, all fish were fasted for the same amount of time (36 h) before the temperature preference test (Killen 2014).

For the test, a single fish was placed in the shuttle-box at static mode, with the cold chamber at 14°C and the warm chamber at 16°C. The fish was left to acclimate overnight with the lights off. The following morning, we turned on the lights at 9:00 and allowed the fish to acclimate for one additional hour. From 10:00 to 18:00, the system was set to dynamic mode, and the fish was allowed to select its preferred temperature. During this time, we also collected data on each individual’s activity (total distance moved).

Core body temperature (*T_b_*) was calculated using the following equation (Killen 2014): *T_b_ = T_a_ + (T_i_ - T_a_)*e^-kt^*. Here, *T_a_* is the current water temperature experienced by the fish, *T_i_* is the previous temperature it experienced, *t* is the time elapsed since experiencing that previous water temperature, and *k* is the rate coefficient for thermal equilibration. The rate coefficient *k* varies with body size and is defined as the instantaneous rate of change in body temperature in relation to the difference between *T_a_* and *T_b_* (Pépino et al. 2015). We calculated *k* using the equation *k* = 3.69\**m*^-0.574^, where *m*
 denotes mass (Stevens and Fry 1974). The final preferred temperature of each individual was calculated as the mean core body temperature experienced during the final two hours of the test when the system was in dynamic mode (Killen 2014).

### Data analysis

R version 3.5.1 was used for all statistical tests (R Core Team 2018), and the ggplot2 package was used for generating figures (Wickham 2009). To test for differences in temperature preference and activity, we used linear models with the following explanatory variables: body mass, population pair (allopatric populations, sympatric population 1, or sympatric population 2; as defined in Table 1), thermal habitat at population of origin (warm or cold), and the interaction between population pair and thermal habitat. Statistical results reported below are the values from the full models for temperature preference and activity. We also tested for a correlation between the activity level and preferred temperature of each individual (Killen 2014).

## RESULTS

There was considerable variation in preferred temperature among individuals, ranging from 8.0°C to 15.8°C (Figure 2). However, there was no difference in mean preferred temperature between sticklebacks collected from warm versus cold habitats (*F_1, 54_*=1.03, *P*=0.31). This finding is unlikely to be due to low statistical power given that the absolute differences in mean preferred temperature between thermal habitats were very small (Figure 2). There was also no evidence for a greater divergence in temperature preference in populations exposed to warm water for more generations (Figure 2). Lastly, temperature preference was not influenced by body mass (*F_1, 54_*=1.29, *P*=0.26), population pair (*F_2,54_*=0.03, *P*=0.97), or the interaction between population pair and thermal habitat (*F_2,54_*=2.87, *P*=0.07).

**Figure 2.**
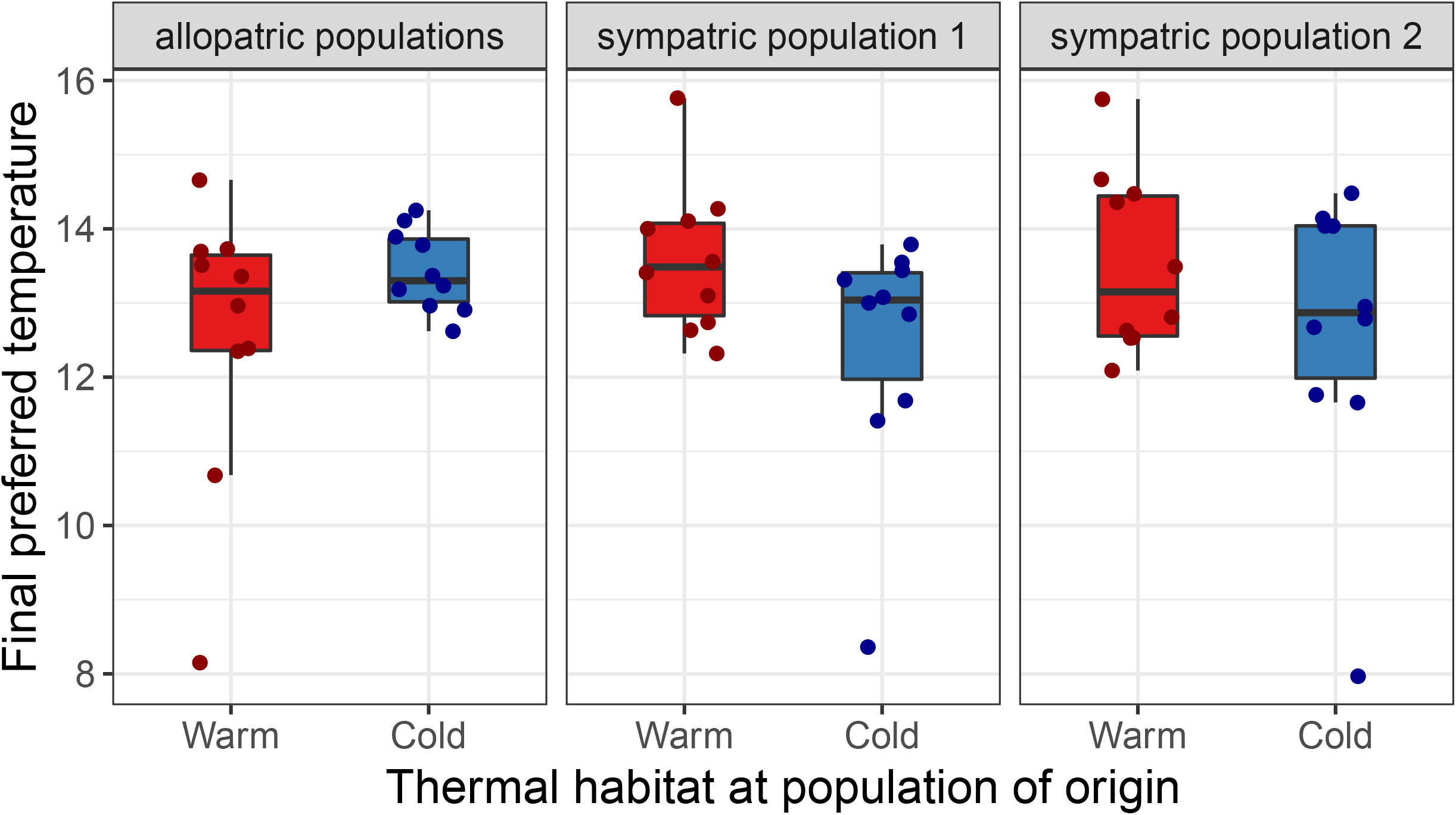
Boxplots of final preferred temperatures (°C) for warm-habitat (red) and cold-habitat (blue) sticklebacks from three population pairs (*n*=10 for sampling location). Filled circles represent individual data points. ‘Allopatric populations’ refers to Grettislaug and Gar?svatn, ‘sympatric population 1’ refers to Áshildarholtsvatn, and ‘sympatric population 2’ refers to Mývatn.

Similarly, we found no differences in activity between fish from warm and cold habitats (*F_1,54_*=0.152, *P*=0.70; Figure 3) or between fish from different population pairs (*F_2, 54_*=0.66, *P*=0.52; Figure 3). Activity also did not depend on body mass (*F_1,54_*=0.003, *P*=0.95) or the interaction between population pair and thermal habitat (*F_2,54_*=2.37, *P*=0.10). Despite a positive trend, there was no statistically significant relationship between individual activity and temperature preference (*t*_58_=1.81, *r*=0.23, *P*=0.076; Figure 4).

**Figure 3.**
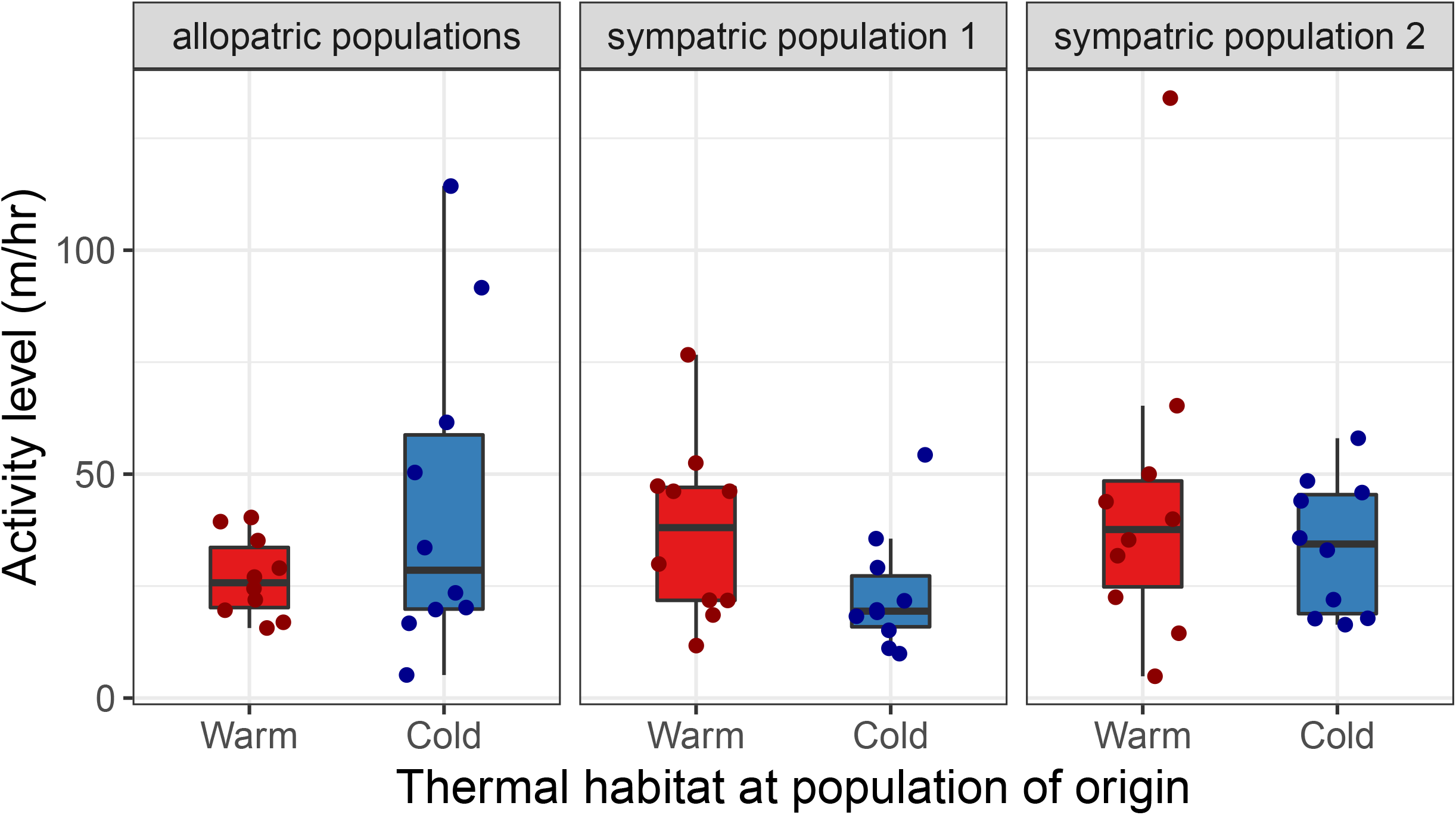
Boxplots of individual activity level (distance travelled in m/hr) of warm-habitat (red) and cold-habitat (blue) sticklebacks from three population pairs (*n*=10 for each sampling location). Filled circles represent individual data points. ‘Allopatric populations’ refers to Grettislaug and Gar?svatn, ‘sympatric population 1’ refers to Áshildarholtsvatn, and ‘sympatric population 2’ refers to Mývatn.

**Figure 4.**
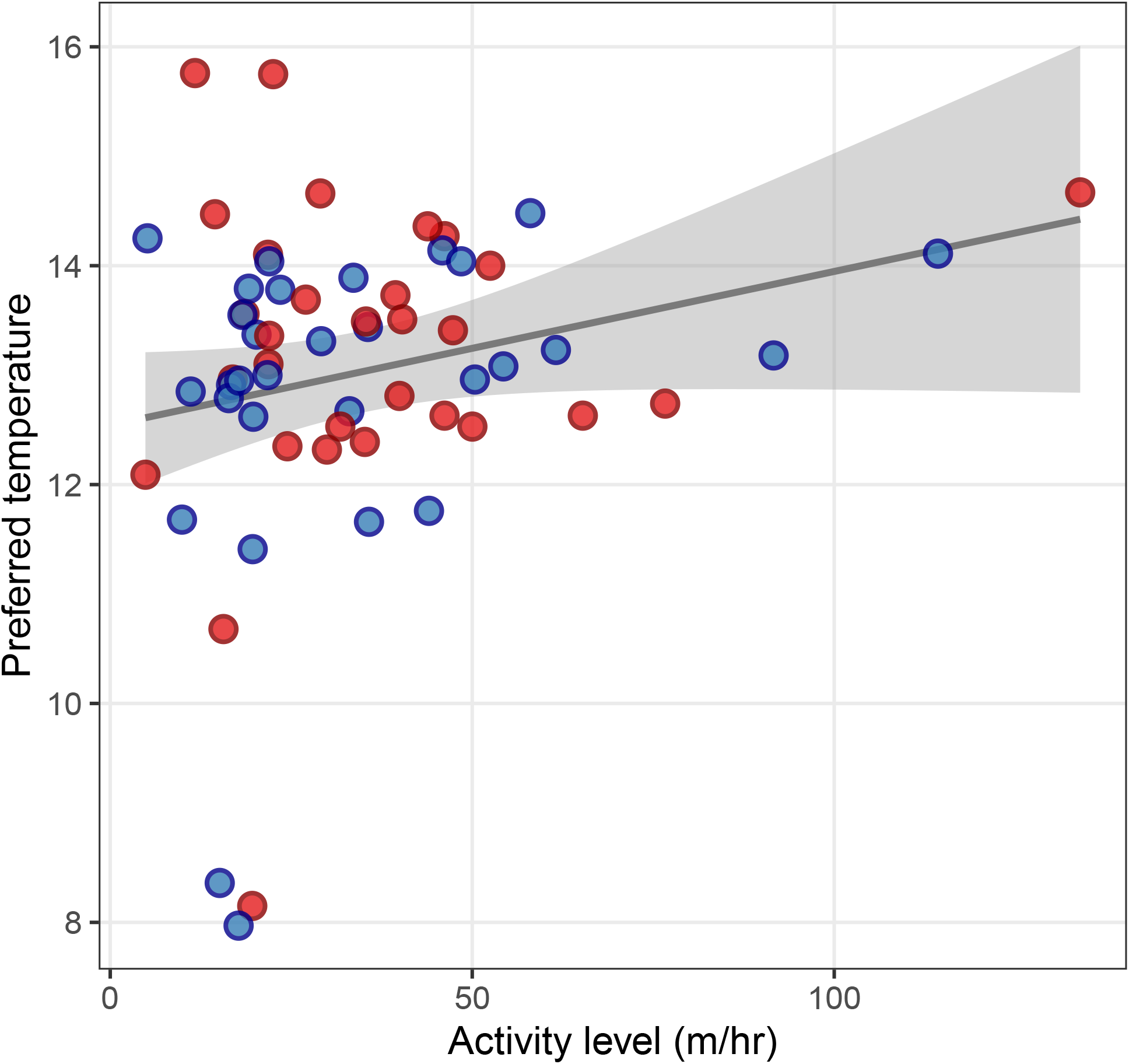
Scatterplot and regression line for the relationship between temperature preference (°C) and activity level (distance travelled in m/hr) of warm-habitat (red) and cold-habitat (blue) sticklebacks. The grey shaded area shows the 95% confidence interval.

## DISCUSSION

Contrary to our hypothesis, fish from warm habitats did not prefer higher water temperatures than fish from cold habitats when acclimated to a common temperature. There was also no evidence for a larger difference in temperature preference between warm- and cold-habitat fish from old populations than from young populations. Average preferred temperature was the same for warm- and cold-habitat fish from allopatric and sympatric populations. Thus, the absence of a difference in sympatric populations is not necessarily due to gene flow constraining adaptation. Lastly, we found no differences in activity between fish from warm and cold habitats, and there was no relationship between activity and temperature preference.

In this study system, populations in the cold habitats likely represent the ancestral populations for those in the warm habitats. Therefore, we considered the average preferred temperature of the cold-habitat fish (13°C) to be the ancestral state. On this basis, our results suggest that temperature preference has not diverged in warm-habitat populations even after being exposed to elevated temperatures for hundreds to thousands of years. This finding is in line with a recent study showing that frogs inhabiting Japanese hot springs prefer cooler water when given the option (Komaki et al. 2020). A similar compensatory response has been observed in the social spider *Stegodyphus dumicola* (Malmos et al. 2021). One explanation for these patterns is that there is a lack of genetic variation in traits associated with temperature preference. Although this may seem surprising, it is consistent with prior work showing low to no heritability of temperature preferences. For example, there is evidence that non-genetic factors may play a more important role in some species. In side-blotched lizards (*Uta stansburiana*), maternal effects but not additive genetic variation influence the offspring’s temperature preference (Paranjpe et al. 2013). Similarly, a study on brown anole lizards (*Anolis sagrei*) found no differences in and no heritability of thermoregulatory behaviour in two populations from contrasting thermal environments (Logan et al. 2018).

Interestingly, the average preferred temperature we observed in this study (13°C) matches the temperature corresponding to peak immune activity in some of these Icelandic populations (Franke et al. 2017, Franke et al. 2019). Previous work using sticklebacks from Lake Mývatn (sympatric population 2) showed that both warm- and cold-habitat sticklebacks had improved acquired immunity at 13°C (Franke et al. 2017, Franke et al. 2019). Such immunity benefits of lower temperatures may also apply to other species, as warmer environments are associated with increased disease risk, virulence, and parasite development (Harvell et al. 2002, Macnab & Barber 2011, Altizer et al. 2013, Franke et al. 2017).

Given that preferred temperature often coincides with optimum temperature for growth, survival, and offspring performance (Jobling 1981, Kellogg & Gift 1983, Diaz et al. 2007, Paranjpe et al. 2013; but see Huey & Bennett 1987, MacLean et al. 2019, Clark et al. 2022), future work should investigate whether temperature preference reflects optimal physiological performance in these populations (Köhler et al. 2011, Artacho et al. 2015, Gilbert & Miles 2017). However, even if performance peaks at a relatively low temperature (as is the case for immune activity), sticklebacks inhabiting warm habitats are clearly able to persist at much higher temperatures during the summer months (Table 1). Hence, we suggest that preferred temperature alone may be a poor indicator of adaptation and evolutionary potential in relation to the thermal environment.

Our results also raise the question of why warm-habitat fish from sympatric populations are found at non-preferred temperatures in the summer, even though they are not restricted by geographic barriers. A plausible explanation is that although only water temperature differed between the two chambers in our experimental set-up, temperature is not the only difference between warm and cold habitats in the wild. Habitat choice is based on many biotic and abiotic factors, which can lead to trade-offs. For example, predation risk may be lower in warm habitats because large piscivorous salmonids are less able to cope with high temperatures (Eliason et al. 2011). Moreover, warm habitats may provide a longer breeding season, leading to a higher reproductive output each annual cycle (Hovel et al. 2016). Previous work has also found differences in prey type availability between these warm and cold habitats in Iceland (Kreiling et al. 2018). Lastly, it is possible that warm-habitat sticklebacks are found at these sites due to social interactions, such as competition or social inertia, rather than active habitat choice (Stodola & Ward 2017, Jacob et al. 2018).

A potential caveat of our study is that all experimental fish were acclimated to a common temperature before the test, which could have led to a plastic response for a similar preferred temperature (MacLean et al. 2019). Nevertheless, we believe this is unlikely, because we still observed substantial among-individual variation in temperature preferences, ranging from 8°C to 16°C (Figure 1). In addition, we have previously shown that metabolic rate differences between warm- and cold-habitat sticklebacks persist after similar long-term acclimation to a common temperature in the laboratory; this is the case for both allopatric and sympatric population pairs (Pilakouta et al. 2020). Prior work in many fish species, including sticklebacks, has also shown that acclimation temperature does not influence final preferred temperature (Røed 1979, Kelsch & Neill 1990, Pérez et al. 2003, Diaz et al. 2007, Habary et al. 2017, Tabin et al. 2018, but see Crawshaw 1975). For example, Røed (1979) showed that sticklebacks acclimated to different temperatures for different amounts of time all showed similar preferred temperatures. Similarly, a study on a coral reef fish (*Chromis viridis*) found that preferences for cooler water persist after prolonged acclimation to high temperatures, due to an inability to acclimate at the level of the metabolic rate (Habary et al. 2017).

In sum, we find no evidence for a divergence in temperature preference between natural populations that have been exposed to contrasting thermal regimes for many generations. Our findings highlight the need to re-evaluate the common assumption that temperature preferences in ectotherms will readily evolve in response to climate change (Kearney et al. 2009, Huey et al. 2012, Gilbert & Miles 2017, Fey et al. 2019). Furthermore, the fact that warm-habitat fish can persist at high temperatures despite a lower temperature preference suggests that preferred temperature alone may be a poor indicator of a population’s adaptive potential to a novel thermal environment.

## ACKNOWLEDGEMENTS

We thank Iain Hill, Tiffany Armstrong, Anna Persson, and Kári Heiðar Árnason for help with fieldwork in Iceland and Joseph Humble for assistance with animal husbandry.

## DATA AVAILABILITY STATEMENT

All relevant data have been uploaded to the Dryad Digital Repository: doi:10.5061/dryad.n2z34tn14.s

## ETHICAL STATEMENT

Our study adheres to the ASAB/ABS Guidelines for the Use of Animals in Research, the institutional guidelines at University of Glasgow, and the legal requirements of the UK Home Office (Project License P89482164).

